# Epicatechin modulates stress-resistance in *C. elegans* via Insulin/IGF-1 signaling pathway

**DOI:** 10.1101/344374

**Authors:** Begoña Ayuda-Durán, Susana González-Manzano, Antonio Miranda-Vizuete, Montserrat Dueñas, Celestino Santos-Buelga, Ana M. González-Paramás

## Abstract

The nematode *Caenorhabditis elegans* has been used to examine the influence of epicatechin (EC), an abundant flavonoid in the human diet, in some stress biomarkers (ROS production, lipid peroxidation and protein carbonylation). Furthermore, the ability of EC to modulate the expression of some key genes in the insulin/IGF-1 signaling pathway (IIS), involved in longevity and oxidative or heat shock stress response, has also been explored. The final aim was to contribute to the elucidation of the mechanisms involved in the biological effects of flavonoids. The results showed that EC-treated wild-type *C. elegans* exhibited increased survival and reduced oxidative damage of biomolecules when submitted to thermal stress. EC treatment led to a moderate elevation in ROS levels, which might activate endogenous mechanisms of defense protecting against oxidative insult. The enhanced stress resistance induced by EC was found to be mediated through the IIS pathway, since assays in *daf-2, age-1, akt-1, akt-2, sgk-1, daf-16, skn-1* and *hsf-1* loss of function mutant strains failed to show any heat-resistant phenotype against thermal stress when treated with EC. Consistently, EC treatment upregulated the expression of some stress resistance associated genes, such as *gst-4, hsp-16.2* and *hsp-70*, which are downstream regulated by the IIS pathway.

## Introduction

Flavan-3-ols, such as epicatechin (EC), catechin (C) and their oligomers, the procyanidins, represent a major class of secondary polyphenolic plant metabolites. Flavan-3-ols are among the most abundant flavonoids in the human diet and are mainly present in fruits, tea, cocoa and red wine. These compounds have been reported to exhibit a range of biochemical and pharmacological activities [1], although their precise mechanisms of action have not been yet elucidated. Traditionally it has been assumed that antioxidant and radical scavenging properties underlay their action mechanism, but currently it is not clear whether other pathways contribute to their overall effect and could be even more important than the radical scavenging properties [2].

Aging is a degenerative process that is receiving increasing attention in recent years. The latest theories suggest that aging is in fact a multifactorial process that is often associated with an increase of oxidative stress leading to cellular damage, as well as by gene mutation due to developmental, genetic and environmental factors [3, 4, 5]. Oxidative stress is an imbalanced state in which excessive quantities of reactive oxygen species (ROS) overcome the endogenous antioxidant capacity of a biological system, leading to an accumulation of oxidative damage in a variety of biomacromolecules, such as enzymes, proteins, DNA, and lipids [6].On the other hand, ROS have been found to be physiologically vital for signal transduction, gene regulation and redox regulation among others, implying that their complete elimination would be harmful [7].

*Caenorhabditis elegans* is a simple multicellular organism that constitutes an excellent model for studying mechanisms of aging because of its short lifespan, fast generation time, good molecular and genomic tools and well-defined genetic pathways [8,9]. Furthermore, *C. elegans* molecular and cellular pathways are strongly conserved in relation to mammals, including humans. Comparison between human and *C. elegans* genomes confirmed that many of human genes and pathways involved in disease development are present in the worm [10]. Thus, the use of *C. elegans* offers promising possibilities for studying the influence of secondary plant compounds like flavonoids on the process of aging and human health [2].

The aging, metabolism and stress resistance processes are regulated by an environmental conserved insulin/IGF-I signaling (IIS) pathway (Fig 1) [3, 11].

**Fig 1.**
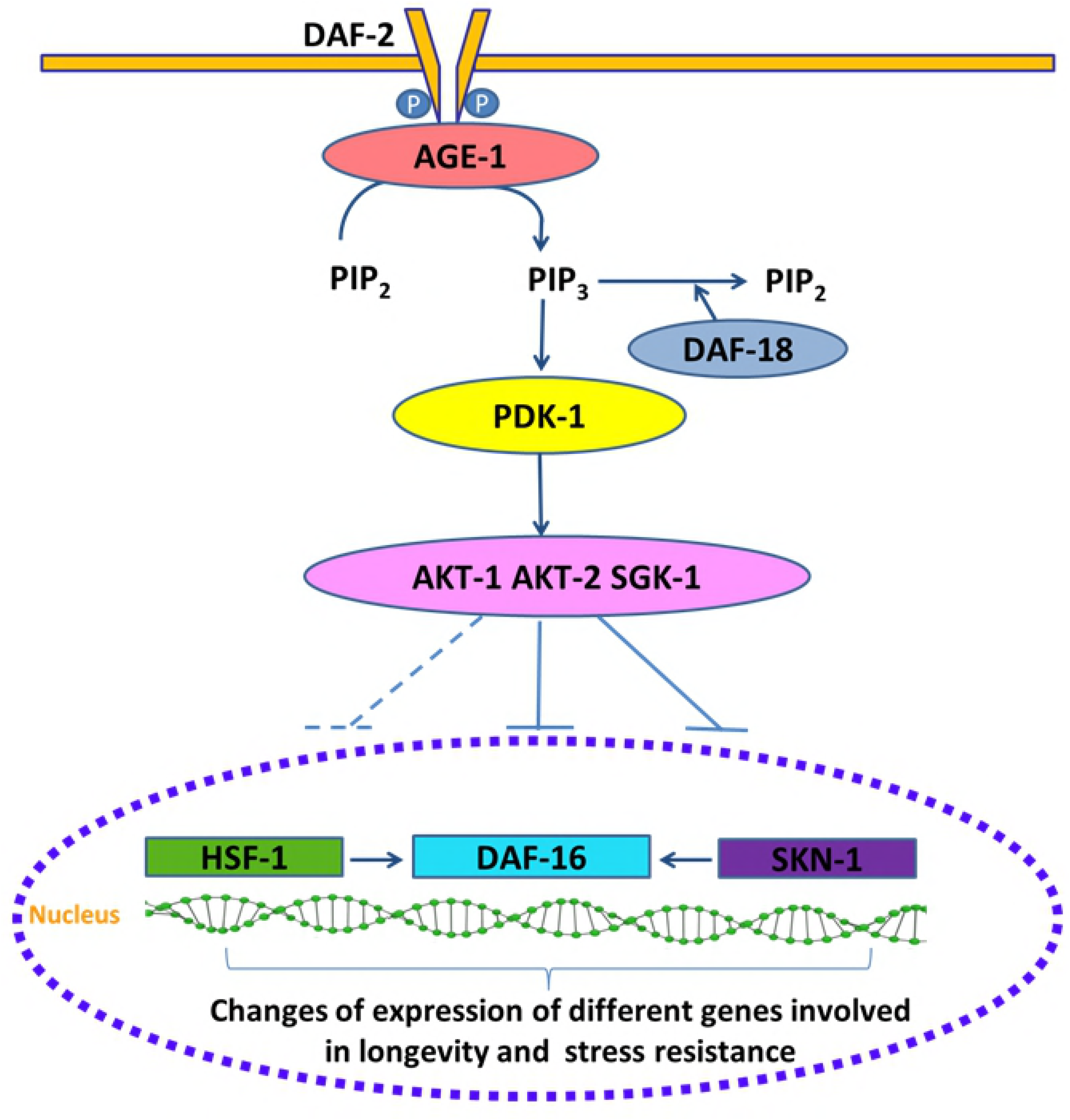
Scheme of *C. elegans* IIS pathway.

Components of this pathway are novel candidate targets, which could provide a powerful entry point for understanding the causes of aging at the molecular level. The IIS pathway consists of DAF-2, a receptor tyrosine kinase that gets phosphorylated upon stimulation by insulin-like peptides (ILPs) and promotes the activation of a phosphatidylinositol 3-kinase signaling cascade that culminates in the phosphorylation and inactivation of DAF-16/FOXO transcription factor by promoting its nucleus-cytosol translocation [12, 13]. The inhibition of the IIS pathway by an increased DAF-18/PTEN activity, stress or reduced DAF-2 activity, leads to nuclear translocation and activation of DAF-16/FOXO, where it changes the expression of various genes. DAF-16/FOXO interacts with other transcription factors such as HSF-1 and SKN-1 that are also affected by DAF-2 [14]. These transcription factors, in turn, regulate the expression of many genes such as catalase *(ctl-1)*, superoxide dismutase-3 *(sod-3*), metallothionein (*mtl-1*), bacterial pathogen defense genes (*lys-7, spp-1*), molecular chaperones, e.g., small heat shock protein-16.2 (*hsp-16.2*) and glutathione *S*-transferase (*gst-4*). All of them key factors that contribute to lifespan, stress tolerance, response to pathogenic bacteria and protein misfolding suppression [15-18].

Therefore, mutations in DAF-2 or any of the other downstream signaling components produce the downregulation or inhibition of IIS signaling in *C. elegans* and cause several cytoprotective phenotypes, such as stress resistance (oxidative stress, thermal stress), increased pathogen resistance and long lifespan [8]. In the case of *daf-2* mutants the lifespan of the animal is increased more than double and the most remarkable issue about these (and many other) long-lived mutants is that they remain young and healthy long after wild type worms are old and decrepit [4]. Previous studies have shown that different phenolic compounds such as acacetin [19], quercetin [20, 21], epicatechin [22], epigallocatechin-3-*O*-gallate (EGCG) [23] or myricetin [24], seem to have an influence in this pathway and/or have the ability to prolong lifespan or attenuate oxidative stress.

In this work, besides the study of the influence of EC in the biochemical changes on wild type *Caenorhabditis elegans*, genetic analyses within a series of worm mutants of the IIS pathway (*daf-2, age-1, daf-16, akt-1, akt-2-; sgk-1, hsf-1, skn-1*) have been carried out in order to evaluate the effects of EC on oxidative resistance. Additionally, the expression of some of these stress resistance associated genes, such as *daf-16, skn-1 hsf-1, hsp-16.2, hsp-70, sod-3 and gst-4* has been determined by quantitative real-time PCR or using transgenic strains expressing fluorescent reporters. The aim of these studies is to gain further insight into the mechanisms involved in the effects of EC in aging.

## Material and methods

### Standards and reagents

(-)-Epicatechin (EC), 2’-7’dichlorofluorescein diacetate (DCFH-DA), ampicillin sodium salt, nistatine, agar, yeast extract, fluorodeoxyuridine (FUdR), phosphate-buffered saline (PBS), cholesterol, Bradford reagent, guanidine hydrochloride (GuHCl), 2,4-dinitrophenylhydrazine (DNPH), malondialdehyde, hexanal, hexenal and 4-HNE were purchased from Sigma-Aldrich (Madrid, Spain). Dimethyl sulfoxide (DMSO) was obtained from Panreac (Barcelona, Spain) and trichloroacetic acid from Fluka Analytical (Madrid, Spain). HPLC grade acetonitrile was from Carlo Erba (Rodano, Italy). Acetic acid was from Merck (Darmstadt, Germany). Fluorescein thiosemicarbazide was from Carbosynth (Berkshire. UK)

### Strains and Maintenance Conditions

The wild type strain N2 and the mutant strains CB1270, *daf-2* (e1370) III; TJ1052, *age-1*(hx546) II; CF1038, *daf-16*(mu86) I; CB1375, *daf-18*(e1375) IV; BQ1, *akt-1*(mg306) V; KQ1323, *akt-2*(tm812) *sgk-1*(ft15) X; PS3551, *hsf-1*(sy441) I; EU1, *skn-1(zu67)* IV*/nT1[unc-?(n754)let-?]* (IV;V); CF1553, *muls84 [(Psod-3::gfp)]*; TJ356, *zIs356 [Pdaf-16::daf-16::gfp; rol-6 (su1006)]* IV; CL2166, *dvIs19 [(Pgst-4::gfp::NLS; rol-6 (su1006)]* III; AM446, *rmIs223 [Phsp70::gfp; rol-6(su1006)]*; CL2070, *dvIs70 [Phsp-16.2::gfp]; rol-6 (su1006)]*, as well as the *E. coli* OP50 bacterial strain were obtained from the *Caenorhabditis* Genetics Center at the University Minnesota (Minneapolis, USA). Worms were routinely propagated at 20 °C on nematode growth medium (NGM) plates with OP50 as a food source.

Synchronization of worm cultures was achieved by treating gravid hermaphrodites with bleach:NaOH 5N (50:50). Eggs are resistant whereas worms are dissolved in the bleach solution. The suspension was shaken with vortex during one min and kept a further minute on rest; this process was repeated five times. The suspension was centrifuged (2 min, 9500 *g*). The pellet containing the eggs was washed six times with an equal volume of buffer M9 (3 g KH_2_PO_4_, 6 g Na_2_HPO_4_, 5 g NaCl, 1 mL 1 M MgSO_4_, H2O to 1 L). Around 100 to 300 μL of the M9 with eggs (depending on eggs concentration) were transferred and incubated on NGM agar plates. When the worms reached the L4 stage they were transferred to new plates with or without EC but also containing FUdR at a concentration of 150 μM to prevent reproduction and progeny overgrowth. The worms were transferred every 2 days to fresh plates with FUdR for the different treatments (with or without EC) until they reached the day of the assay. Epicatechin solution (200 mM) in DMSO was added to the nematode growth medium during its preparation to get a 200 μM final concentration on the plates. Control plates were also prepared without the flavonoid but containing the same volume of DMSO (0.1% DMSO, v/v).

In order to evaluate if the developmental stage of the worm had an influence, the different assays were carried out at different stages of development as described below.

### Stress Assays

Oxidative stress in worms was induced by subjecting the animals to 35°C heat-shock treatment. Worms were incubated on OP50 plates with or without EC until days 10 and 17 of adulthood for wild type worms, and days 2 and 9 of adulthood in mutant worms. Then they were transferred with a platinum wire to agar plates (∅ 35 mm, 20 worms per plate) and switched to 35 °C for 6 or 8 h. The time was decided depending on the thermotolerance of the specific strain used in the assay. After that time, dead and alive nematodes were counted. Assays were performed with approximately 100 nematodes per treatment. In all mutant assays, in addition to the mutant control a parallel control using wild type worms was also included. In all cases, three independent experiments were performed. The relative rates of survival of worms after being subjected to thermal stress were expressed in relation to the untreated controls.

### Determination of Reactive Oxygen Species (ROS)

The accumulation of ROS was evaluated periodically every two days from the 2^nd^ day to the 17^th^ day of adulthood in worms cultivated in presence and absence of EC. The cellular ROS were quantified by the dichlorofluorescein assay [25]. Briefly, the worms were individually transferred to a well of a 96-well plate containing 75 μL of PBS and then exposed or not to thermal stress (2 h at 35 °C), after which 25 μL of DFCH-DA 150 µM solution in ethanol was added to each well. The acetate groups of DFCH-DA were removed in worm cells, and the released DFCH is oxidized by intracellular ROS to yield the fluorescent dye DCF. The fluorescence from each well was measured immediately after incorporation of the reagent and every 10 minutes for 30 minutes, using 485 and 535 nm as excitation and emission wavelengths, respectively. Recording of the DCF fluorescence intensity with time in single worms was used as an index of the individual intracellular levels of ROS. Five independent experiments were performed per treatment, and for each experiment ROS measurements were made in at least 24 individual worms. The measurements were performed in a microplate reader (FLUOstar Omega, BMG labtceh).

### Worm homogenates

Worms were grown on NMG medium until the 10^th^ and 17^th^ day of adulthood. Then, they were subjected to thermal stress for 5 h at 35 °C and subsequently, for each assay, animals from two plates (∅ 100 mm) were collected to a flask and resuspended in M9 buffer. Suspensions were centrifuged (12,000 *g*, 5 min), and the worm pellet was washed with PBST (PBS + 0.01% Tween 20) twice and finally with PBS. The remaining pellet was transferred to an Eppendorf tube, resuspended in 1000 mL of PBS, and kept at −20 °C. Next, samples were stirred (Genius 3 vortex) and sonicated once during 60 s and twice for 30 s in a Cell Disruptor (Microson XL2000 100) to obtain a homogenate. For each treatment three independent experiments were performed, and in each experiment the measurements of the different variables were made in triplicate using three different worm homogenates. The protein content was determined according to the Bradford method after digestion of the homogenate [26]. The carbonylated proteins and lipid peroxidation products were further normalized to protein content to correct for differences in biomass of the different homogenates.

### Determination of lipid peroxidation products

Lipid peroxidation products were analyzed by HPLC after derivatization with 2,4-dinitrophenylhydrazine (DNPH) based on the method described by Andreoli et al. [27]. Proteins were removed from worm homogenates (350 µL) by adding 350 µL of 20% (v/v) trichloroacetic acid; 100 µL of butylhydroxytoluene 10 mM dissolved in methanol was also added in order to protect the lipids. After a 15 min incubation at 4 °C, samples were centrifuged at 10,000 *g* for 10 min at 4 °C. The supernatant was mixed with 100 µL of 10 mM DNPH in 2M HCl and incubated for 60 min at room temperature. The mixture was extracted three times with 400 µL of chloroform and 3 pieces of molecular sieves were added to the organic phase for 30 min in order to remove possible remains of aqueous phase. The organic phase was collected and concentrated to dryness and finally resuspended in 80 µL of acetic acid 0.2%: acetonitrile (62:38, v/v) and injected in the HPLC system. The column was a Waters Spherisorb S3 ODS-2 C8, 3 µm (4.6 × 150 mm) and the solvents were: (A) 0.2% acetic acid, and (B) acetonitrile. The elution gradient established was: isocratic 38% B for 10 min, 38% to 75% B over 10 min, 75% to 80% B over 20 min at a flow rate of 0.6 mL/min. Malondialdehyde, 4-hydroxynonenal and cis-hexenal were used as lipid peroxidation markers. Double online detection was carried out in a DAD using 310 nm and 380 nm as preferred wavelengths, and in a mass spectrometer for compound confirmation. MS detection was performed in negative ion mode in an equipment provided by an APCI source and a triple quadrupole-ion trap mass analyzer. The APCI temperature was set at 450 °C. Lipid peroxidation products were quantified from their chromatographic peaks recorded in the DAD by comparison with calibration curves obtained by injection of increasing concentrations of malondialdehyde (310 nm), hexenal and 4-hydroxynonenal (HNE) (380 nm).

### Determination of carbonylated proteins

Carbonylated proteins were determined by a direct reaction of protein carbonyls with fluorescein thiosemicarbazide (FTC) [28] and measured in a fluorescent semi-microplate assay. A 50 µL of 0.2 mM of FTC was added to 50 µL of homogenate and kept overnight. Proteins were precipitated by adding 400 µL 20% trichloroacetic acid and centrifuged 10, 000 *g* 4 °C 10 min. Afterwards, the precipitate was cleaned three times with 1 mL acetone, stirred (Genius vortex) and centrifuged for 10 min at 10,000 *g* 4 °C. The precipitates were dried and finally solubilized with 50 µL of 6M guanidine hydrochloride (GuHCl). The samples were diluted with 450 µL Hepes buffer 0.1 M pH 7 (1.38 g NaH_2_PO_4_.H_2_O dissolved in 100 mL of water) and measured using 100 µL per well in triplicate in a fluorescent reader with excitation at 485 nm and emission at 520 nm. Nanomol of FTC-reacted carbonyls were calculated using a standard curve generated from the readings of various concentrations of FTC prepared in a medium similar to the one used in the samples. The levels of protein carbonyls in the homogenates were expressed as nmol/mg worm protein calculated by the Bradford method.

### RT-qPCR assays

Adult worms were treated with or without 200 μM of EC for 4 days. The worms were collected with M9 buffer, centrifuged at 10,000 *g* 1 min, and the pellet dissolved in 300 µL of M9. Total RNA was extracted using RNAspin Mini RNA Isolation Kit (GE Healthcare). In order to maximize cell breakage, in the first stage of the extraction 10 stainless steel beads (2 mm) were added. The mixture was vortex shaken vigorously and further homogenized in a Thermo Savant FastPrep 120 Cell Disrupter System with a speed of 5.5 m/s and run time duration of 10 s five times. cDNA was produced with High Capacity cDNA Reverse Transcription Kits (Applied Biosystems) using a 2 µg of total RNA per reaction. The expression of mRNA was assessed by quantitative real-time PCR, using SYBR green as the detection method. The gene expression data were analyzed using the comparative 2-ΔΔCt method with act-1 as the normalizer [29]. Nine independent experiments were performed. The following gene-specific primers were used: *act-1* CCAGGAATTGCTGATCGTATG (F) and GGAGAGGGAAGCGAGGATAG (R), *skn-1* AGTGTCGGCGTTCCAGATTTC (F) and GTCGACGAATCTTGCGAATCA (R), *daf-16* CCAGACGGAAGGCTTAAACT (F) and ATTCGCATGAAACGAGAATG (R), and *hsf-1* GAAATGTTTTGCCGCATTTT (F) and CCTTGGGACAGTGGAGTCAT (R).

### Fluorescence quantification and visualization

Synchronized L1 larvae expressing an inducible green fluorescent protein (GFP) reporter for *gst-4, hsp-16.2, hsp-70, sod-*3 and *daf-16* genes were grown on NMG medium in the presence or absence of EC until the day of the assay, when they were submitted or not to thermally-induced oxidative stress (35 °C, 1h). The precise day of assay was defined when a higher intensity of the fluorescence was observed after carrying out a screening with the different strains throughout the life of the worm. If no increase in the fluorescence was observed, young (day 2 ^th^ of adulthood) and older adult worms (day 9^th^ of adulthood) were exposed to the heat shock. In the cases of *hsp-16.2* and *hsp-70* reporter strains, worms were then allowed to recover in their normal environment at 20 °C for 2h or 3h, respectively before pictures were taken. The expression of *gst-4, hsp-16.2, hsp-70, sod-3* was measured by quantifying the fluorescence of the GFP reporter. To analyze the subcellular localization of DAF-16::GFP, worms were classified as diffuse cytoplasmic, intermediate cytoplasmic/nuclear and strong nuclear translocation. Approximately 35 randomly selected worms for each experiment were mounted in a 5 µL drop of 10 mM levamisole (except for DAF-16::GFP in 2% sodium azide) on a 3% agarose pad covered with a coverslip. All fluorescence determinations were done in an Olympus BX61 fluorescence microscope equipped with a filter set (excitation 470±20 mn, emission 500±20 nm) and a DP72 digital camera coupled to CellSens Software for image acquisition and analysis. ImageJ software was used to quantify fluorescence intensity. Three independent experiments were performed per assay and reporter strain.

### Statistical Analysis

The statistical analyses were performed using the PC software package SPSS (version 23.0; SPSS Inc., Chicago). ANOVA was applied for multiple comparisons of values to determine possible significant differences between treated and control groups. To analyze survival to thermal stress, contingency tables were performed and Statistical significance was calculated using the Chi Square Test. In every analysis, significant differences were statistically considered at the level of *p* < 0.05.

## Results and Discussion

### Effects of epicatechin (EC) in stress resistance

In a previous work, the effects of catechin, epicatechin, 3′-O-methylepicatechin and 4′-O-methylepicatechin in *C. elegans* stress resistance were evaluated [30]. All the assayed catechins enhanced the resistance of the worm against both thermal and chemically-induced oxidative stress in early stages of development (worms at 1^st^ and 6th day of adulthood), with relatively greater protective effects in older (6^th^ day) than in young worms. Specifically, a significant enhancement in survival was observed following thermal stress in the EC-treated nematodes (200 μM); in the first day of adult the average proportion of living worms was 78.6% in control assay and 97.6% in treated worms while, in the 6^th^ day of adulthood, survival rate was 89,2% in treated worms compared to 56,2% in untreated animals.

In the present work, the influence of EC in worm resistance to thermal stress was evaluated in more aged animals (10^th^ and 17^th^ day of adulthood), in order to know if the developmental stage of the animals and/ or a longer exposure time to EC further influenced the resistance against this type of stress. As shown in Fig 2, the treatment with EC resulted in a significant increase in the survival of nematodes subjected to thermal stress (8h, 35 °C). At day 10, the survival of stressed animals increased from 29.9% in controls to 47.7% in worms treated with EC. Likewise, the treatment with EC increased the survival rates at day 17 from 40% in controls to 55% in worms treated with EC. These results suggest that the protective effect of EC against oxidative stress is not increased in more aged worms, as previously concluded [30]. Nevertheless, caution must be observed when interpreting these data as this aged population (10^th^ and 17^th^ days of adulthood) represents the more aging resistant phenotypes, a circumstance that might provide a special relevance to the increase in the percentage of survival induced by EC in older individuals.

**Fig 2.**
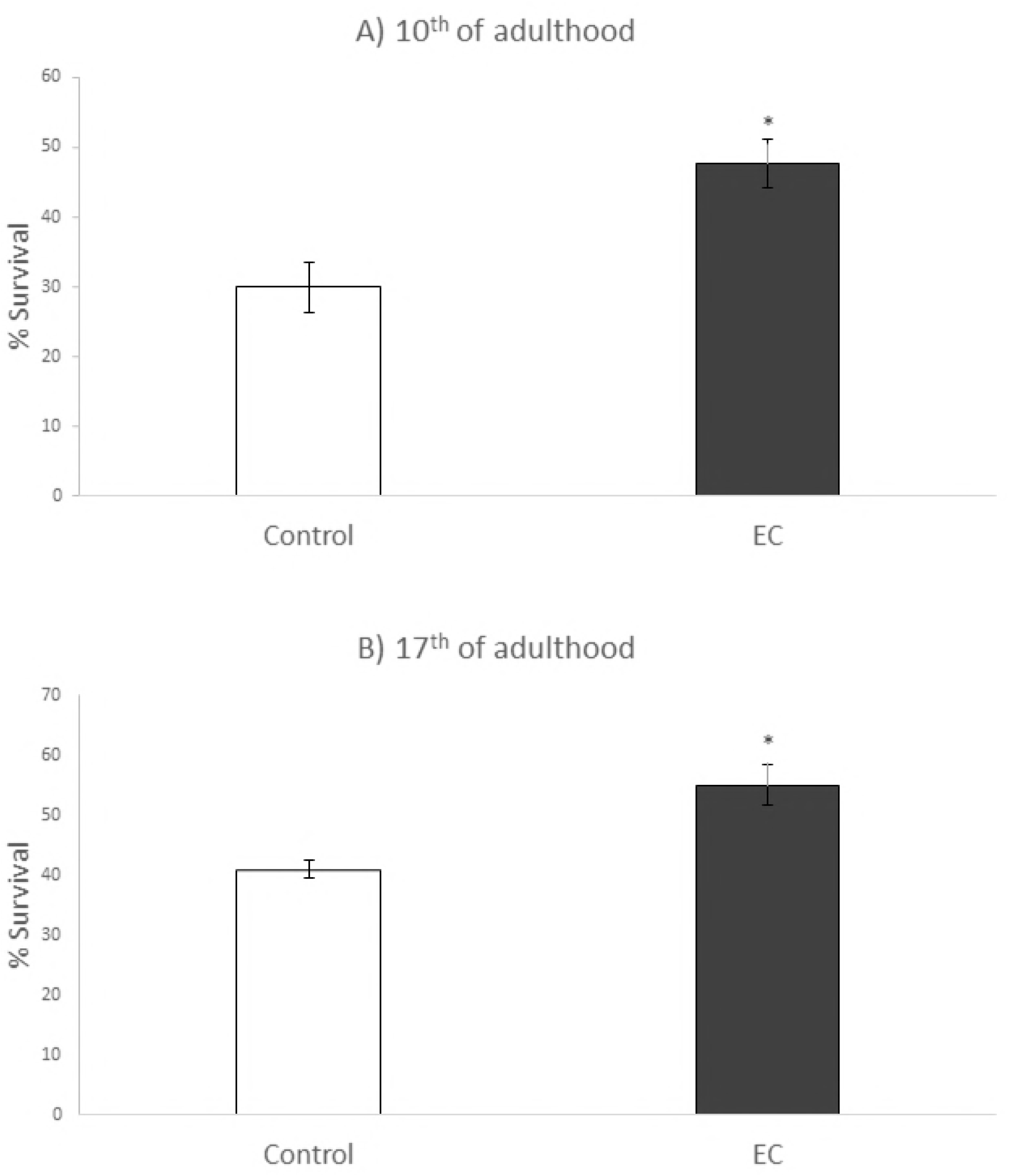
Percentages of survival following thermal stress (35 °C, 8h) applied at days 10^th^ (A) and 17^th^ of adulthood (B) in N2 wild type *C. elegans* strain not treated (controls) and treated with EC (200 µM in the culture media). Three independent experiments were performed. The results are presented as the mean values±SD. Statistical significance was calculated using the Chi Square Test. The differences were considered significant at *(*p*<0.05).

### Effects of EC in intracellular ROS levels

Intracellular ROS were determined in *C. elegans* grown in NGM media with and without EC (200 µM) and exposed or not to thermal stress (35 °C, 2h). ROS assessment was performed every two or three days throughout the life of the worms and the obtained results are shown in Fig 3.

**Fig 3.**
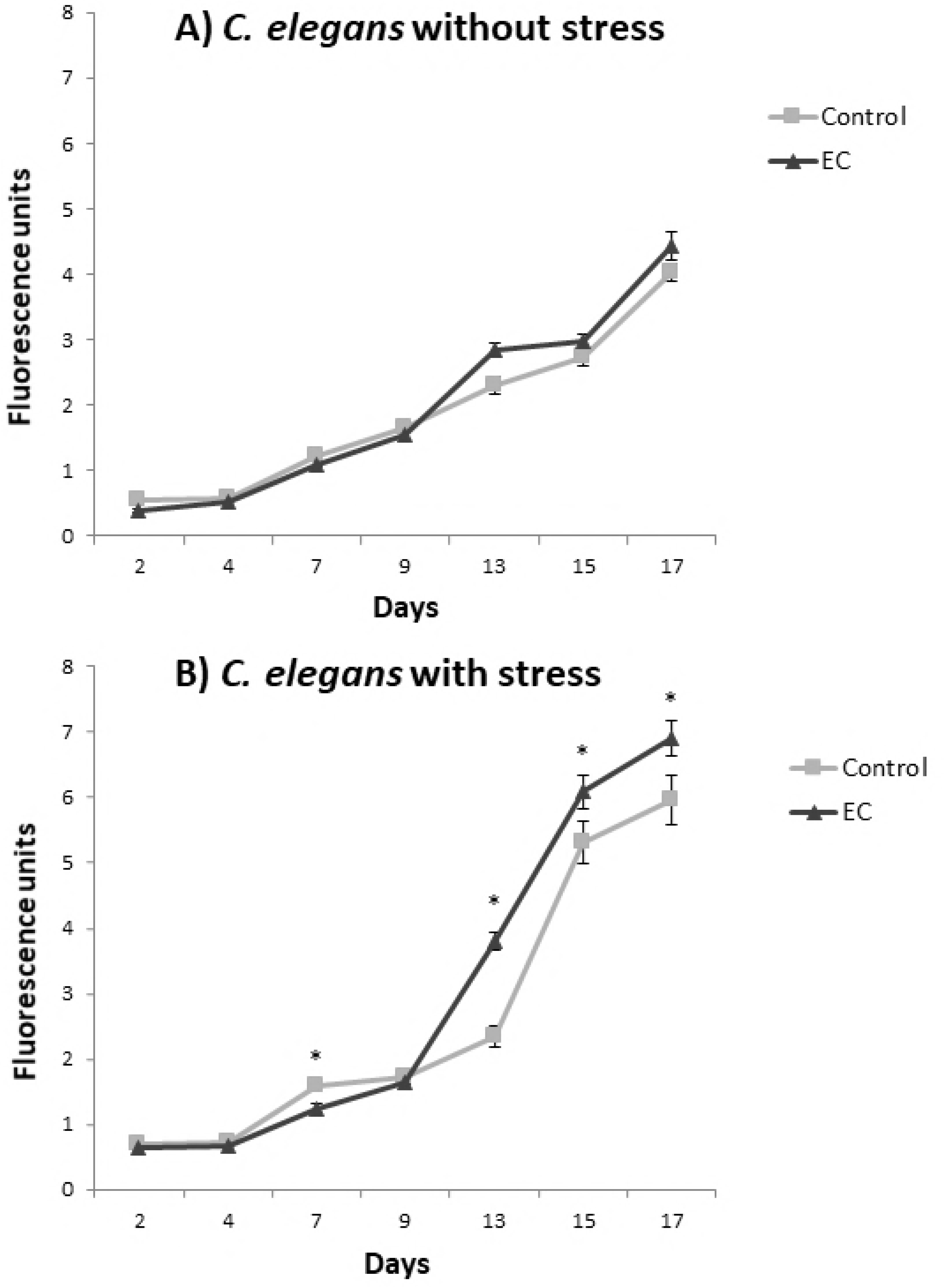
Levels of intracellular ROS in *C. elegans* subjected (B) or not (A) to thermal stress (35 °C, 2h) grown in the absence (controls) or presence of EC (200 μM in the culture media). ROS levels were evaluated at different stages of development throughout the entire life of the worm. Five independent experiments were performed. The results are presented as the mean values ± SEM. Statistical significance was calculated using one-way analysis of variance ANOVA. The differences were considered significant at *(*p*<0.05).

As expected, a progressive increase in ROS levels was produced as the animals grows older and higher ROS levels were found in thermally stressed animals than in those not subjected to stress. Regarding the effect of EC, a different behavior was observed between younger and older individuals. Thus, up to day 9 of adulthood, similar or slightly lower ROS levels were determined in animals treated with EC than in non-treated controls. This observation was in agreement with previous studies where *C. elegans* was grown with and without EC up to the sixth day of adulthood [22]. However, from day 9 onwards this trend was inverted and higher ROS values were determined in worms grown in the presence of EC than in their corresponding controls, either submitted or not to thermal stress. In previous studies on the influence of EC in *C. elegans* longevity [30], an increase in the survival rate was observed in the worms treated with EC from day 14^th^ onwards, which approximately coincides with the time point where ROS levels become higher in the individuals treated with EC in both populations in the assays now performed (Fig 3).

The physiological effects of ROS levels within an organism remains unresolved. According to the free radical theory of aging [31], the cause of aging is the accumulation of molecular damage due to the production of toxic reactive oxygen species during cellular respiration. Nevertheless, although it is clear that oxidative damage increases with age, studies both in invertebrate (worms and flies) or mammals (mice) have suggested that oxidative stress may not be the only cause of aging or at least not according to the classical conception [5, 32, 33]. Indeed, an increasing number of studies seem to contradict the free radical theory, including studies carried out in *C. elegans* were longer lifespan was found in worms with higher concentrations of ROS. Lee et al, [34] showed that the mild increase in ROS levels induced by the inhibition of respiration in *C. elegans* stimulates HIF-1 to activate gene expression and promote longevity. These same authors observed that low paraquat levels, an oxygen free radical generating compound, increased worm lifespan significantly whereas higher concentrations of paraquat decreased it in a dose-dependent manner. Similarly, Heidler et al, [35] observed that exposure to high concentrations of juglone, another superoxide-generating compound, led to premature worm death but low concentrations prolonged life. In that study, lifespan extension was associated with an increased expression of small heat-shock protein HSP-16.2, enhanced glutathione levels and nuclear translocation of DAF-16. Based on the observations above, Van Raamsdonk and Hekimi [36] proposed that *C. elegans* lifespan resulted from a balance between pro-survival ROS-mediated signaling and ROS toxicity. According to those authors, superoxide was not a simply toxic byproduct of metabolism, but it is involved in a type of ROS-mediated signaling that can result in increased longevity.

More recently, Meng et al, [37] studied the differential responses to oxidative stress in young and old individuals using *C. elegans* and human fibroblasts. They proposed a new concept called “Redox-stress Response Capacity (RRC)”, according to which cells or organisms are capable of generating dynamic redox responses to activate cellular signaling and to maintain cellular homeostasis. This response would be higher in young individuals generating more ROS and activating signaling pathways and with a better ability to degrade damaged proteins by up-regulating chaperones. That explanation might give an answer to our and others observations regarding the differential effects of EC on ROS production and *C. elegans* survival depending on worm life stage.

Taken together, and in agreement with what has been proposed by other authors [5, 35, 38], the results obtained herein seem to reinforce the emerging idea that mild increase in ROS levels may have beneficial effects. This might involve different mechanisms, such as induction in the expression of protective cellular pathways, activation of repair mechanisms or changes in respiration.

### Oxidative damage: Protein carbonylation and products of lipid peroxidation

In order to evaluate whether the treatment with EC had an influence on the level of oxidative damage in C*. elegans*, carbonylated proteins and lipid peroxidation products were determined in wild type worms grown in the presence and absence of EC (200 µM) and subjected to thermal stress at 10^th^ and 17^th^ day of adulthood.

Carbonylated proteins are commonly used as a biomarker of protein oxidation in cells and tissues and high levels of them have been related to loss of cell viability. The oxidation status of proteins was quantified after the reaction of the carbonyl groups with fluorescein-thiosemicarbazide (FTC) adapting the method proposed by Chaudhuri et al, [28] to *C. elegans*. The results were expressed as nmol of carbonylated proteins by mg of worm protein. As shown in Fig 4A.1 and 4A.2) a slight decrease was observed in the levels of protein carbonylation in worms treated with EC both at days 10^th^ and 17^th^. Although the differences were not significant (p> 0.05), the levels of carbonylated proteins were never higher in the worms treated with EC with respect to untreated animals. This observation suggested that exposure to EC did not lead to an increase in the oxidative damage despite enhanced ROS levels were determined in treated worms than in controls (Fig 3).

**Fig 4.**
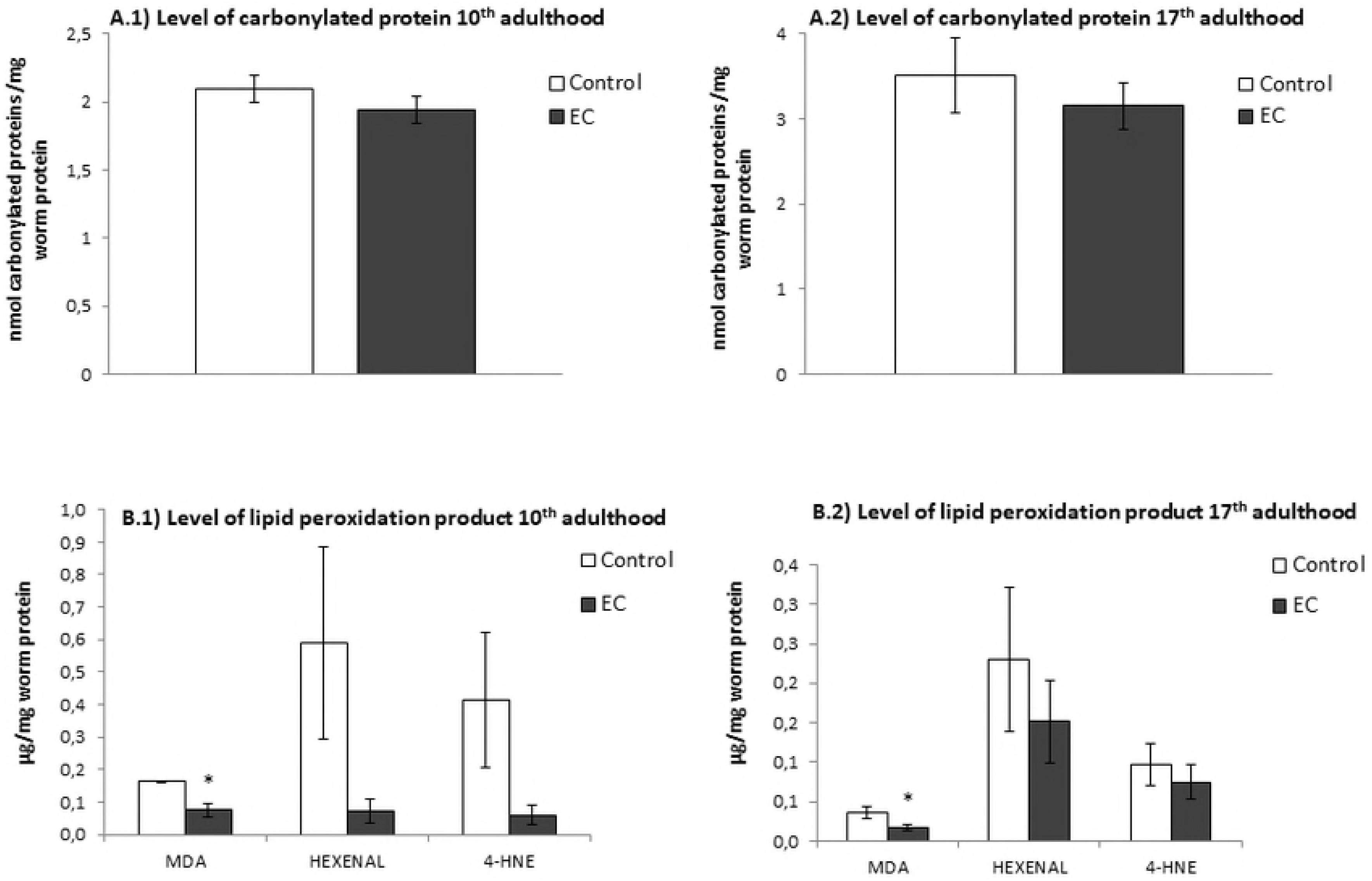
Levels of (A) carbonylated proteins and (B) lipid peroxidation products after cultivation of *C. elegans* in the absence (controls) and presence of EC (200 µM) and subjected to thermal stress. The results were obtained at days 10^th^ (A.1 and B.1) and 17^th^ (A.2 and B.2) of worm adulthood. Three independent experiments were performed. The results are presented as the mean values ± SEM. Statistical significance was calculated using one-way analysis of variance (ANOVA). The differences were considered significant at *(*p*<0.05).

High ROS levels may attack polyunsaturated fatty acids in membrane and free lipids leading to oxidative lipid degradation. Some common products of this process are malondialdehyde (MDA), 4-hydroxynonenal (4-HNE) and cis-hexenal, which have been used as lipid peroxidation markers in the present study. In Fig 4 (B.1 and B.2) it can be observed that a descent was produced in the levels of these peroxidation products in the worms treated with EC with respect to untreated animals in both days of the assay, although only the decline of MDA was significant. Thus, as for carbonylated proteins, the increased ROS levels determined in the worms treated with EC (Fig 3) did not result in an increase in lipid peroxidation, as evaluated by the analyzed markers.

Lipid peroxidation can cause loss of membrane integrity and subsequent cell death. Thereby, the observed decrease in lipid peroxidation might explain the increased survival rate and longer life duration in worms treated with EC submitted to thermal stress. This could indicate that chronic exposure to this flavonoid confers protection against oxidative damage, an effect that would be especially evident in later stages of the life of *C. elegans*, as deduced from the observations made in the longevity and thermal stress resistance assays.

In view of these findings, it could be suggested that the moderate increase ROS levels provoked by the treatment with EC in *C. elegans* leads to a compensatory response, inducing some endogenous antioxidant and other protection mechanisms, which would result in greater resistance to oxidative stress. As proposed by Xiong et al. [38] for similar results in worms treated with EGCG, this type of response could be explained by a specific process of hormesis, the mitohormesis. While hormesis refers to a biphasic dose response to an environmental agent or chemical agent characterized by a low dose adaptive beneficial effect and a high dose toxic effect, the mitohormesis is the hormetic reaction in response to mitochondrial ROS, by which a high but sub-lethal level of free radical production that can stimulate resistance to ROS damage and increase longevity [39]. A support to this assumption can be found in the observations made by Lapointe and Hekimi and Lapointe et al. [5,40] in long-lived *Mclk1*^+/−^ mice, with a dysfunction in the activity of CLK-1/MCLK1, a mitochondrial enzyme necessary for ubiquinone synthesis, which showed a significant attenuation in the rate of development of oxidative biomarkers of aging (protein carbonylation, lipid peroxidation and 8-OHdG as a biomarker of DNA damage) despite they exhibited a substantial increase in oxidative stress. Reduced activity of CLK-1/MCLK1 has also been shown to prolong average and maximum lifespan in *C. elegans* [40].

### Influence of EC on genes involved in oxidative stress resistance

The idea that flavonoids do not act in the organism only as conventional antioxidants but could also modulate multiple cellular pathways is currently gaining strength [41]. The IIS pathway contributes to longevity and oxidative or heat shock stress response and it encompasses highly conserved components from nematodes to mammals, including humans [18]. Some authors have reported that several classes of flavonoids seem to influence this pathway [42-44]. However, although there are many works about the beneficial effects of different flavan-3-ols and flavan-3-ol-rich extracts in different organisms including humans, the molecular mechanisms involved in such effects have not been sufficiently studied.

In the present work, those molecular mechanisms have been explored by checking the ability of EC to modulate the stress resistance in mutant worms for different genes of the IIS pathway and genes that are relevant to stress resistance. The premise of these assays was that EC treatment would not increase the survival of nematodes lacking specific genes that are required for the protection against oxidative damage induced by submitting worms to thermal stress. The stress resistance has been studied in mutant worms at 2^nd^ and 9^th^ day of adulthood, in order to check whether the results could be different according to the developmental stage. Thus, young adults in reproductive age and older adults in post-reproductive age were chosen. Furthermore, the effect of EC on the expression of some of these genes by RT-qPCR in EC-treated worms grown under non-stress conditions and after thermal stress was also investigated.

DAF-2 is the *C. elegans* homologue for the insulin/IGF-1 receptor. Activation of DAF-2 leads to phosphorylation and cytoplasmic sequestration of the DAF-16 transcription factor via AGE-1, PDK-1, AKT-1, AKT-2, and SGK-1 kinases [2]. Herein, the influence of EC on the resistance to thermally-induced stress was checked in *age-1, akt-1, akt-2; sgk-1* and *daf-2* loss of function mutant strains and we found that the treatment with the flavonoid did not lead to significant enhancement in the stress resistance in any of these mutant strains (Fig 5). This result suggests that those genes could be required to explain the mechanisms involved in the effects of the studied flavonoid on improving the resistance against thermal/oxidative stress in *C. elegans* and also that the resistance to stress mediated by EC involves the IIS pathway. Nevertheless, it is also necessary to take into account that these mutants are long-lived and already more resistant to stress than wild type worms, which might mask a possible increase in the survival of the stressed animals produced by EC.

**Fig 5.**
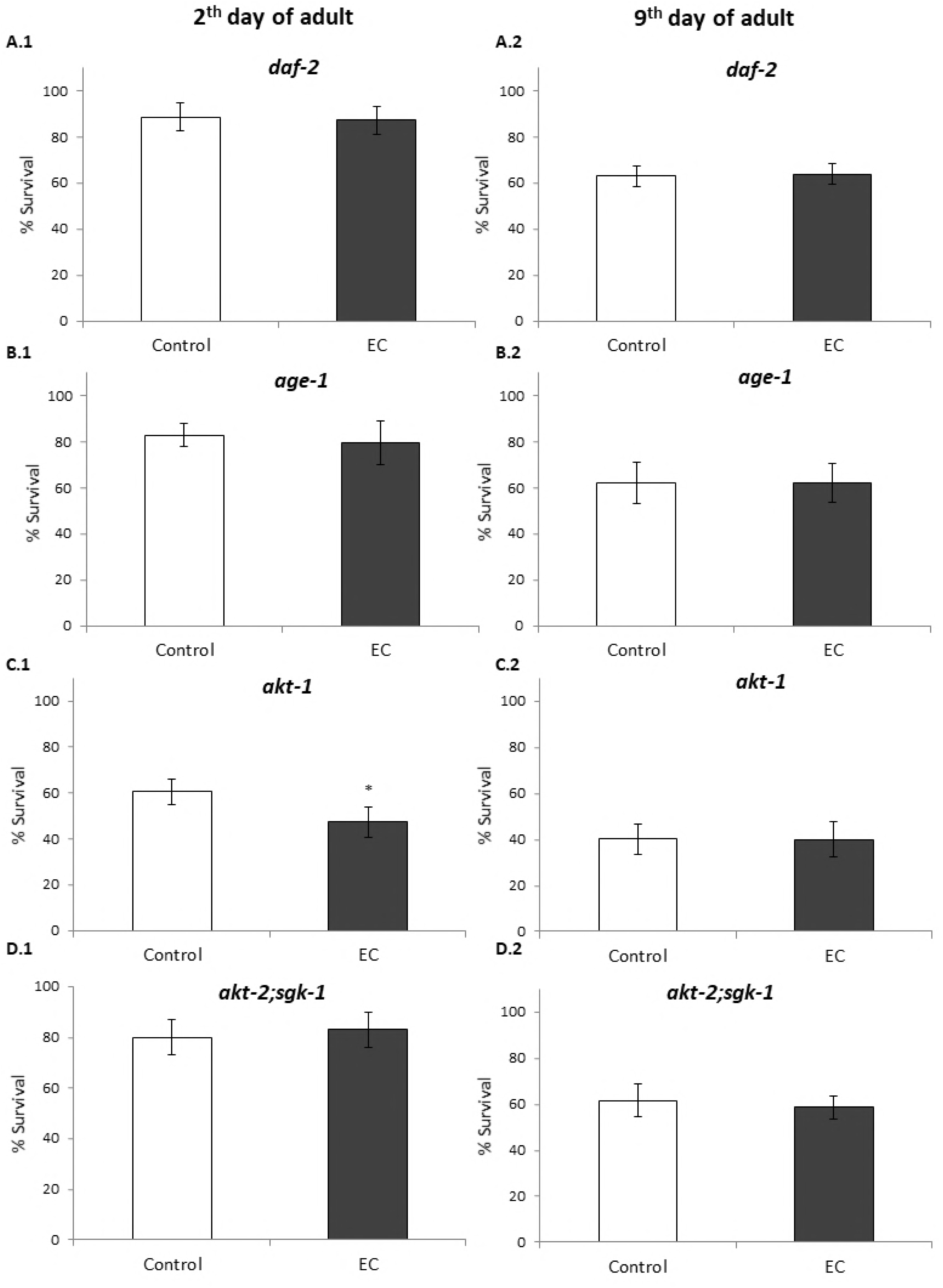
Percentages of survival following thermal stress applied at days 2^nd^ and 9^th^ of adulthood in different long-lived *C. elegans* mutants from the IIS pathway cultivated in the absence (controls) and presence of EC (200 μM) in the culture media. Three independent experiments were performed. The results are presented as the mean values±SD. Statistical significance was calculated using the Chi Square Test. The differences were considered significant at \**(p*<0.05).

Proper regulation of IIS is crucial for the protection of *C. elegans* from both external and internal stresses [14]. The key downstream transcription factors of IIS pathway that contribute to longevity and regulate the resistance to a variety of stress include DAF-16/FOXO, HSF-1 and SKN-1 [14]. Thus, we examined the oxidative stress resistance of loss-of-function *daf-16*, *hsf-1* and *skn-1* mutant worms treated with EC. The results showed that treatment with EC did not increase the survival of these mutants (Fig 6), suggesting that these genes are required for EC-mediated enhanced thermal stress resistance in *C. elegans*. Similar results were obtained for both young adults (day 2) and older worms (day 9).

**Fig 6.**
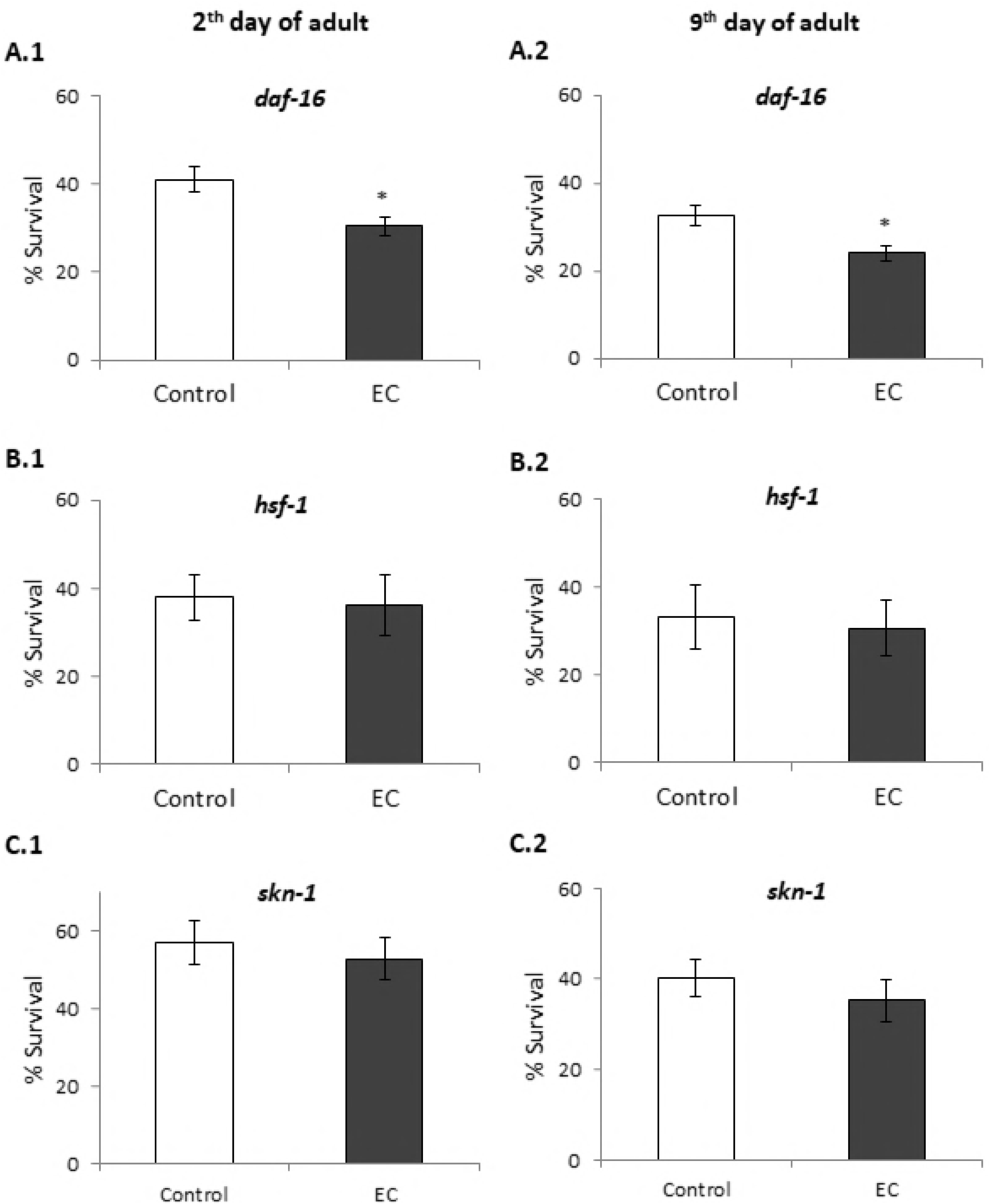
Percentage of survival following thermal stress applied at days 2nd (A, C and D) and 9th (B, D and F) of adulthood in *daf-16(mu86)*, *hsf-1(sy441)* and *skn-1(zu67)* mutants cultivated in the absence (controls) and presence of EC (200 μM) in the culture media. Three independent experiments were performed. The results are presented as the mean values±SD. Statistical significance was calculated using the Chi Square Test. D The differences were considered significant at \**(p*<0.05).

Contrary to our observations, Saul et al. [45] found that 200 µM of catechin significantly prolonged the lifespan in *age-1* and *daf-16* mutants, indicating that AGE-1 and DAF-16 would not be required for the life-extending effect of this flavan-3-ol. However, no significant lifespan extension was observed in *akt-2* mutants, suggesting that AKT-2 was at least partly involved in the catechin mediated longevity. Those authors concluded that the IIS-pathway was not required for the life extending effect of catechin and that the results obtained for AKT-2 could be explained because of a possible AKT-2 function independent of IIS pathway. On the contrary, Cai et al. [46] reported that the lifespan extension effect of the flavonol icariside II was dependent on the IIS pathway, since *daf-16* and *daf-2* loss-of-function mutants failed to show any lifespan extension upon treatment with this compound.

DAF-16, a FOXO-family transcription factor, influences the rate of aging in response to insulin/insulin-like growth factor (IGF-1) signalling by upregulating a wide variety of genes including cellular stress-response, lifespan, antimicrobial and metabolic genes [18]. As above discussed, the treatment with EC did not enhance resistance to thermal stress of *daf-16(mu86)* mutants worms, either at days 2 or 9 of adulthood (Fig 6 A and B), pointing to DAF-16 being involved in EC activity. In order to obtain further support to this assumption, the effect of EC on *daf-16* expression in wild-type *C. elegans* under normal growth conditions and after thermal stress exposure was examined by RT-qPCR. It was found that *daf-16* mRNA levels were enhanced in worms grown in the presence of 200 µM of epicatechin, both subjected and not to thermal stress, although this increase was only significant in worms grown under non-stressed conditions (Fig 7A and 7B). These results support the idea of DAF-16 playing a key role in the effects produced by EC in worms.

**Fig 7.**
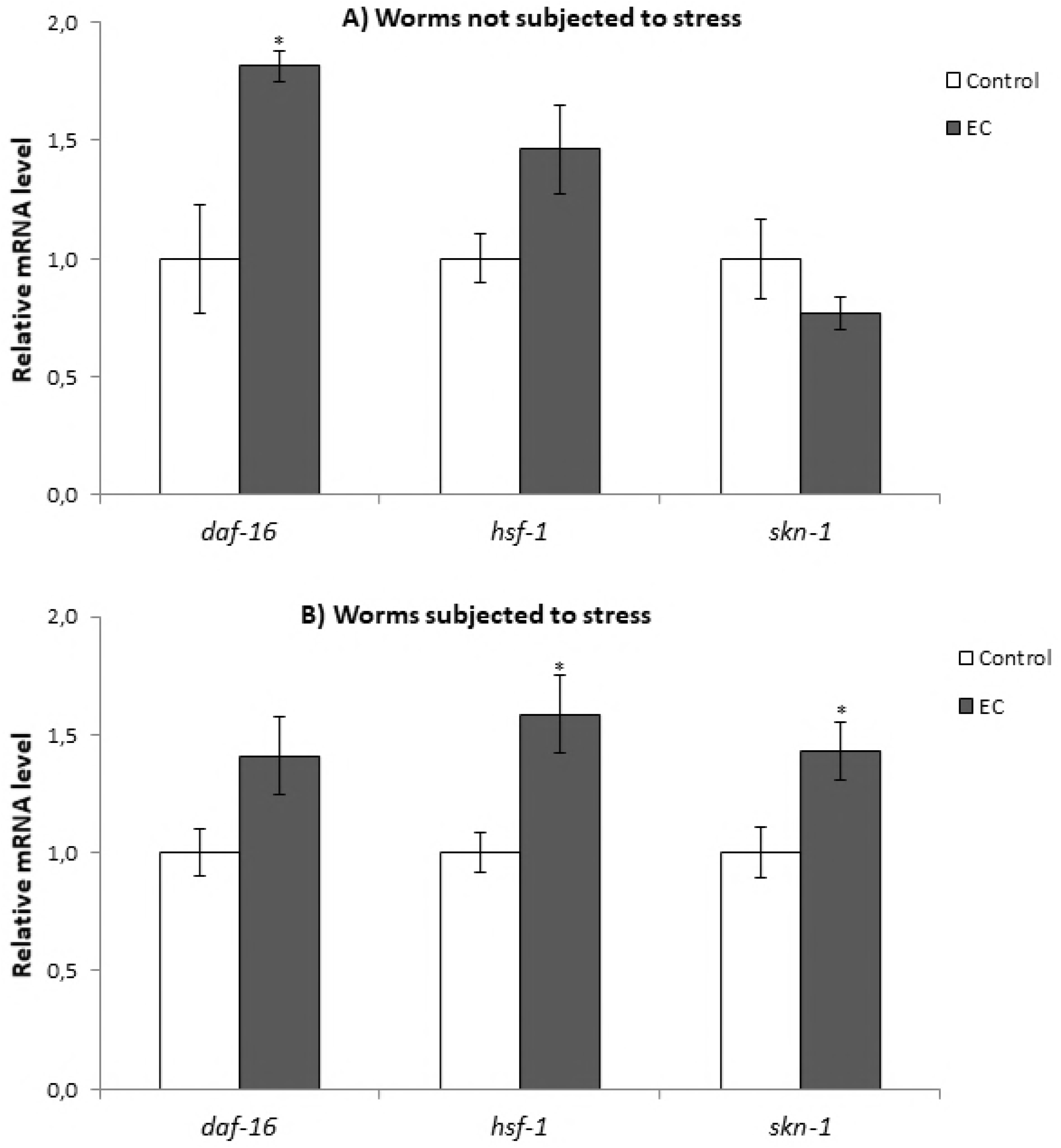
Effect of EC on the expression of *daf-16*, *hsf-1* and *skn-1* genes in wild-type *C. elegans* cultivated in the absence (controls) and presence of EC (200 μM) in the culture media grown under non-stressed conditions (A) or after subjecting them to thermal stress (B). The expression level was determined by RT-qPCR; *act-1* was used as an internal control. Nine independent experiments were performed. The results are presented as the mean values ± SEM. Statistical significance was calculated using by one-way analysis of variance ANOVA The differences were considered significant at (\**p*<0.05).

HSF-1 is a transcription factor that regulates heat shock response and also has an influence in aging [17]. As for *daf-16*, the expression of *hsf-1* was quantified in wild type worms under normal growth conditions and after thermal stress. The results showed an increase in the expression of this transcription factor in both conditions although the increase was only significant only in thermal stress conditions (Fig 7A and B). These results, together with the fact that EC did not increase the resistance to thermal stress of *hsf-1* mutants (Fig 6C and D), could indicate that *hsf-1* is also involved in the effects produced by EC in the worms. Similar observations were made regarding SKN-1 homologue of Nrf-2 transcription factor, which regulates lifespan and oxidative stress response by mobilizing the conserved phase 2 detoxification response [16]. In this case, RT-qPCR experiments showed that EC significantly increased the expression of *skn-1* under stress but not in normal growth conditions (Fig 7A and B). These results, together with the survival assays in which no significant increase was observed in the survival of EC-treated *skn-1(zu67)* mutants compared to control worms (Fig 6E and F), also suggested the involvement of SKN-1 in the effects of EC. Altogether, these results indicated that the improvement in stress resistance produced by EC involves the IIS pathway by regulating the expression of *daf-16*, *hsf-1* and *skn-1* genes independently of the worm age.

In line with the results obtained herein, higher resistance to oxidative stress and increased lifespan was found in *C. elegans* treated with a flavonoid-enriched cocoa powder that contained catechin, epicatechin and procyanidins, which was explained to be mediated by the IIS pathway and sirtuin proteins [42]. Similar studies with chlorogenic acid also concluded that this polyphenol activates the transcription factors DAF-16, HSF-1, SKN-1 and HIF-1, although not SIR-2.1 [47]. By contrast, Saul et al [48] found that the forkhead transcription factor DAF-16 was not essential for quercetin effects on longevity and stress resistance. The same group showed that quercetin-mediated lifespan extension was neither a caloric restriction mimetic effect nor a sirtuin (sir-2.1) dependent process, but it was modulated by four genes: *age-1, daf-2, unc-43* and *sek-1*, identified as a likely mode of action [20]. These observations might indicate that different mechanisms of action could be involved in the effects on longevity and stress resistance induced by different polyphenols.

### Effect of epicatechin on DAF-16 subcellular localization and expression of GST-4, HSP-16.2, HSP-70 and SOD-3

In order to delve into the molecular mechanisms involved in the stress and lifespan modulation, the effect of EC on the expression of the specific cellular stress response genes *sod-3* (superoxide dismutase), *gst-4* (glutathione-S-transferase), *hsp-16.2* and *hsp-70* (heat-shock proteins) was explored. SOD-3 is an antioxidant enzyme that protects against oxidative stress by catalysing the removal of superoxide. The gene *sod-3* is thought to be a direct target of DAF-16 as the *sod-3* promoter contains consensus DAF-16/FOXO-binding elements (DBEs) [49]. GST-4 enzyme is involved in the Phase II detoxification pathway, playing an important role in resistance to oxidative stress; its expression is mediated by SKN-1 [50]. Heat shock proteins (HSP) are induced in response to thermal and other environmental stresses. The expression of *hsp* genes is mainly regulated by heat shock transcription factor (HSF-1), which is also influenced by the IIS pathway in *C. elegans* [2]. For this study, transgenic strains expressing GFP under the control of *gst-4, sod-3, hsp-16.2* and *hsp-70* promoters were used. Also, a transgenic strain expressing a fusion protein DAF-16::GFP was used to examine whether EC treatment activated DAF-16 nuclear translocation under normal and stress conditions. EC (200 µM) was found to significantly enhance the expression levels of GST-4, HSP-16.2 and HSP-70, whereas no differences existed in the expression of SOD-3 (Fig 8). GFP expression levels were determined under non-stressed conditions for *gst-4* and *sod-3*, reporters while for *hsp-16.2* and *hsp-70* reporters, worms had to be previously subjected to a heat shock (35 °C, 1h) and further let to recover at 20 °C for 2h (*hsp-16.2*) or 3h (*hsp-70*). For *hsp-16.2* and *hsp-70* reporter strains fluorescence was hardly detected before heat stress and no differences between the control and EC-treated worms were observed (Fig S1). Regarding DAF-16, EC treatment failed to induce DAF-16::GFP nuclear translocation respect to the control under both in unstressed or under stress conditions (Fig 9). As a short thermal stress (35 °C, 1h) of the DAF-16::GFP reporter strain is enough to provoke DAF-16 nuclear translocation, it is difficult to observe possible differences induced by the treatment with EC.

**Fig 8.**
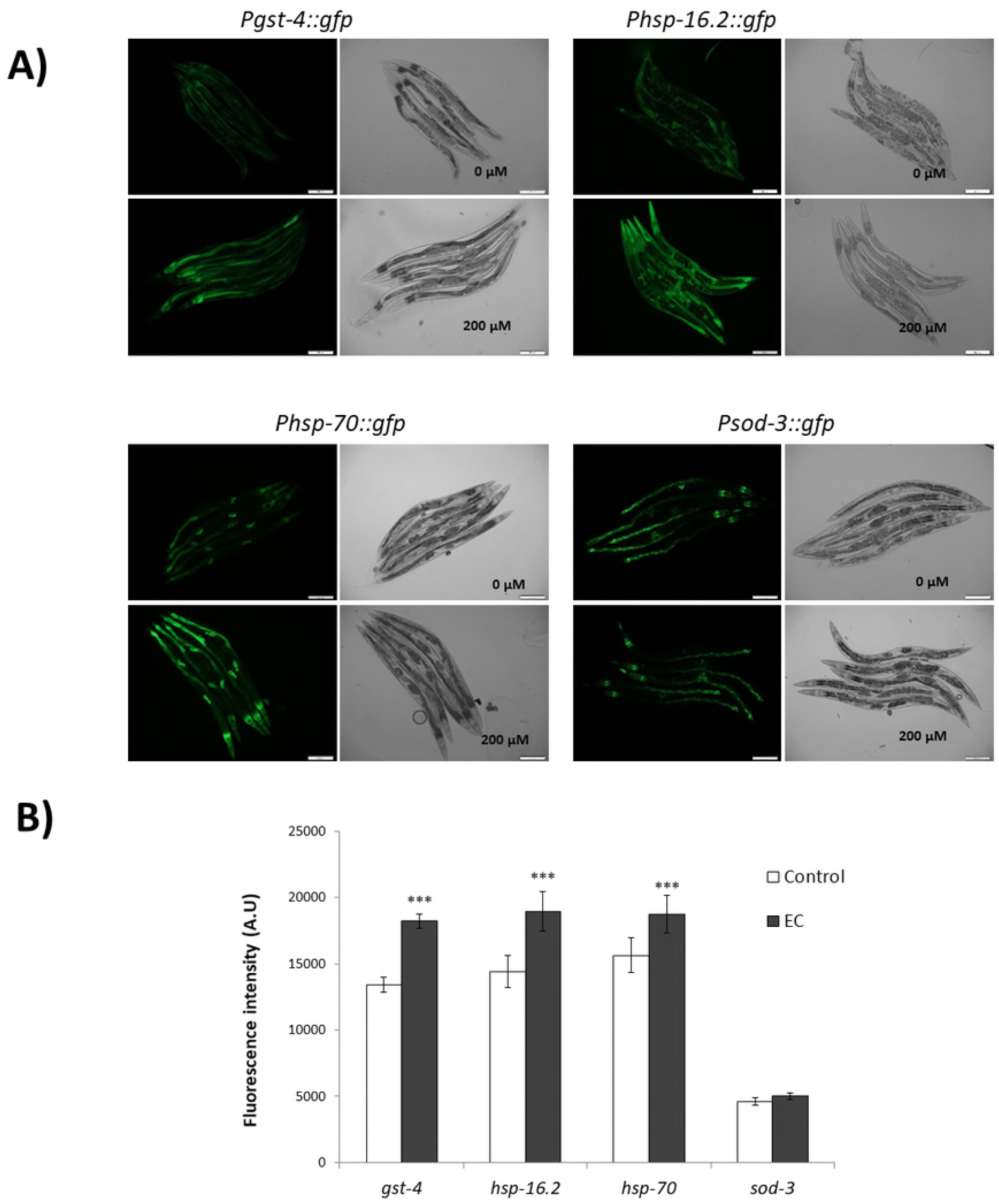
Effect of EC on the expression of GST-4, SOD-3, HSP-16.2 and HSP-70 in *C.elegans*. Age-syncronized L1 transgenic worms of *Pgst-4::gfp*, *Psod-3::gfp*, *Phsp-16.2::gfp* and *Phsp-70::gfp* reporter strains were cultivated in the absence (controls) and presence of EC (200 μM) in the culture media. **A)** Representative fluorescence images of control and EC-treated worm strains stress response. **B)** Relative fluorescence intensities of transgenic worms. Total GFP fluorescence of each whole worm was quantified using Image J sofware. Three independent experiments were performed. The results are presented as the mean values ± SEM. Approximately 35 ramdomly selected worms from each set of experiments were examined. Differences compared with the control (0 µM, 0.1% DMSO) were considered statistically significant at *p*<0.05 (*) and *p*<0.01 (**) and *p*<0.001 (***) by one-way ANOVA.

**Fig 9.**
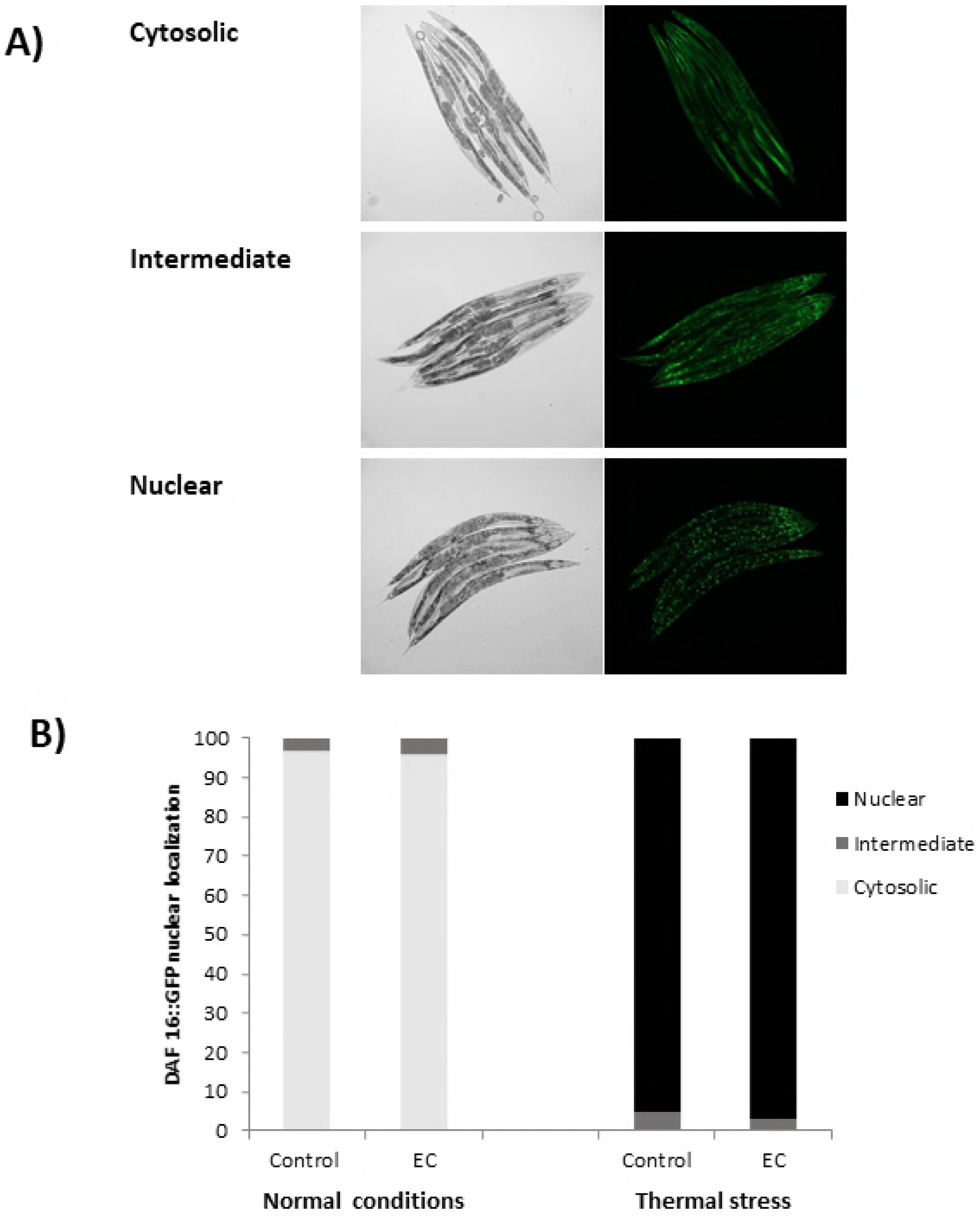
Effect of EC on DAF-16::GFP nuclear localization. Transgenic worms expressing the fusion protein DAF-16::GFP were cultivated in the absence (controls) and presence of EC (200 µM) and evaluated at 2^nd^ day of adulthood. DAF-16::GFP subcellular distribution was classified as cytosolic, intermediate and nuclear.

As previously discussed, EC treatment did not increase oxidative stress resistance in *daf-16(mu86)* mutant nematodes, but it produced an increase in *daf-16* mRNA expression in wild type worms, suggesting that EC protected against thermal stress in a DAF-16-dependent manner. Besides EC treatment did not increase SOD-3::GFP expression, which is coherent with the results obtained in a previous study of our group [22], where no increase in the activity of SOD was found after treatment of the worms with EC. Bonomo et al. [44] obtained similar results in worms treated with a polyphenols-rich extract of Açaí, with no observation of an increase in oxidative resistance in *daf-16(mu86)* mutant worms, as well as no increase in DAF-16 nuclear localization and sod-3 expression under normal conditions. According to those authors [44], this might be explained as the polyphenols extract would lead DAF-16 protein to increase its transcriptional activity but not its concentration, thus DAF-16 activation in the nucleus leading to the upregulation of specific genes other than *sod-3*. In fact, they also observed that the extract increased the expression of genes *ctl-1* and *gst-*7 in a DAF-16 dependent manner. In the same way, our results suggested that the treatment with EC produced a more important effect in other DAF-16 target genes, like *hsp-16.2* and *hsp-70* that encode heat shock proteins.

The GFP expression in some reporter strains studied (*Phsp-16.2::gfp, Psod-3::gfp and Pdaf-16::daf-16::gfp*) was also investigated in older worms (9^th^ day of adulthood) in order to know if the mode of action of EC changed depending on the age of the worm. Similar results were obtained as in younger worms, with no differences in DAF-16 nuclear translocation (Fig. S2) and in the expression of SOD-3 between control and EC-treated worms being observed (Fig. S3). However, the increase expression of HSP-16.2 after thermal stress compared to controls was more accentuated in older worms (Fig. S3s). This observation is relevant, because heat shock proteins levels decrease in aged worms leading to an increase of unfolded proteins, so that worms become more sensitive to stress, finally increasing mortality [43].

Our results identify a significant increase of *gst-4* expression, the loss of resistance to thermal stress in *skn-1(zu67)* mutants and increased *skn-1* expression in worms treated with EC, suggesting that EC could be modulating the Nrf2/SKN-1 pathway. The transcription factor SKN-1 is the ortholog of the mammalian Nrf protein, wich induces the expression of phase-II detoxifying enzymes and antioxidant proteins, such as SOD, GST, glutathione peroxidase (GPO) or NAD(P)H:quinone oxidoreductase (NQO-1) [2, 16]. This control is mediated through an antioxidant response elements (ARE) in the promoter region of genes encoding phase II enzymes and antioxidant components. Several additonal ARE-containing genes were predicted to be direct SKN-1 targets, such as GST-4 (gluthatione transferase-4), which acts conjugating the reduced form of gluthatione (GSH) to a variety of toxic substrates including damaged lipids and proteins, thereby decreasing their activity and making them more water soluble favouring their removal [50, 51].The increased SKN-1 activity could explain the decrease of peroxidated lipids and carbonylated proteins in worms treated with EC with respect to untreated animals. In a previous study, our group also showed that the treatment with EC produced a significant increase in the levels of GSH in *C. elegans* with respect to non treated worms [22]. Similar observations were made in assays carried out on astrocytes [52] and HepG2 cells [53], where the treatment with EC activated Nrf2 and increased GSH levels. Furthermore, it is also known that the Nrf-2-ARE pathway is activated by reactive oxygen species [54]. Thus, the moderate increase in ROS levels observed in worms treated with EC (Fig 3) could lead to the activation of this pahtway, ultimately inducing endogenous antioxidant protection and confering a great protection against oxidative damage.

Tullet et al. [16, 55] proposed that the effects of SKN-1 on resistance to oxidative stress and longevity can be dissociated with SKN-1 being required for resistance to oxidative stress but not for the increased lifespan resulting from overexpression of DAF-16. On the other hand, DAF-16 overexpression rescues the short lifespan of *skn-1* mutants but not their hypersensitivity to oxidative stress. This dual function could explain the effects of EC in *C. elegans* observed herein, where EC-treated worms showed improved resistance to thermal stress but not increased mean lifespan.

High levels of HSP promote longevity and are also a predictor of the ability to withstand thermal stress [23, 56]. Hsu et al. suggested that HSF-1 and DAF-16 together activate the expression of specific genes, including genes encoding HSP, which in turn promote longevity [17]. HSP act as molecular chaperones and proteases by preventing the accumulation of aggregated proteins in response to heat and other forms of stress. This activity may prevent oxidized or otherwise damaged proteins from aggregating before they can be refolded or degradated [17]. The results obtained in the present study showed that EC upregulated HSP-16.2 and HSP-70 in *C. elegans*, which might explain why EC significantly increased the survival of *C. elegans* under heat stress and maximun lifespan. Other authors have also related the improvement in lisfespan and increase of thermal stress resistance in *C. elegans* induced by different polyphenols with the capacity to upregulate *hsp* and other genes associated to stress resistance [19, 23, 57].

### Conclusions

Our results suggest a protection of EC against oxidative damage, as evaluated from worm survival and the levels of lipid peroxidation products and protein carbonylation as biomarkers. EC treatment induces a moderate elevation in ROS levels, which might lead to a compensatory response, increasing endogenous mechanisms of protection that would result in prolonged maximum lifespan and greater resistance to oxidative stress. In addition, stress resistance tests revealed that the heat-resistant phenotype against thermal stress was absent in *daf-2, age-1, akt-1, akt-2, sgk-1, daf-16, skn-1 and hsf-1* mutants. Thus, these protective effects could be mediated through regulation of the insulin/IGF-1 signalling pathway, where DAF-16 acts a central regulator and together with HSF-1 and SKN-1 transcription factors control a wide variety of downstream genes with diverse functions that act in stress response and lifespan modulation. In particular, it has been shown that EC could upregulate the expression of GST-4, HSP-16.2 and HSP-70. Overall, the observations of this study indicated that the effects of EC in stress resistance are achieved by the regulation of the expression of different genes of the IIS pathway independently of the worm age.

## Acknowledgments

This work was funded by MINECO (Spanish National Project AGL2015-64522-C2) (BFU2015-64408-P) and FEDER-Interreg España-Portugal Programme (Project ref. 0377_IBERPHENOL_6_E). B.A-D is recipient of PhD fellowships from the Junta de Castilla y Leon (Orden EDU/310/2015). The authors are thankful to Marta Rodríguez-Romero and José Antonio Mora-Lorca for providing the necessary help with the assays of RT-qPCR and fluorescence microscopy, respectively.

## Author Contributions

**Conceptualization:** Ana M. González-Paramás, Celestino Santos-Buelga

**Formal analysis:** Begoña Ayuda-Durán, Susana González-Manzano, Montserrat Dueñas

**Funding acquisition:** Ana M. González-Paramás, Celestino Santos-Buelga

**Methodology:** Begoña Ayuda-Durán, Susana González-Manzano, Antonio Miranda-Vizuete

**Project administration:** Ana M. González-Paramás, Celestino Santos-Buelga

**Resources:** Ana M. González-Paramás, Celestino Santos-Buelga, Antonio Miranda-Vizuete

**Supervision:** Ana M. González-Paramás, Celestino Santos-Buelga, Antonio Miranda-Vizuete

**Writing – original draft:** Begoña Ayuda-Durán, Celestino Santos-Buelga

**Writing – review & editing:** Begoña Ayuda-Durán, Celestino Santos-Buelga, Ana M. González-Paramás, Antonio Miranda-Vizuete, Susana González-Manzano

## Supporting information

**S1 Fig. Effect of EC on the expression of HSP-16.2 and HSP-70 in *C. elegans***. Age-synchronized L1 transgenic worms expressing *Phsp-16.2::gfp* and *Phsp-70::gfp* transgenes were cultivated in the absence (controls) and presence of EC (200 μM) in the culture media. Relative GFP fluorescence intensities in transgenic **A)** *Phsp-16.2::gfp* and **B)** *Phsp-70::gfp* worms were quantified under normal growth conditions and after subjecting worms to thermal stress to 35 °C for 1h. Total GFP fluorescence of each whole worm was quantified using Image J software. Three independent experiments were performed. The results are presented as the mean values ± SEM. Approximately 35 randomly selected worms from each set of experiments were examined. Differences compared with the control (0 µM, 0.1% DMSO) were considered statistically significant at p<0.05 (*) and p<0.01 (**) and p<0.001 (***) by one-way ANOVA.

**S2 Fig. Effect of EC on DAF-16::GFP nuclear localization**. Transgenic worms expressing the DAF-16::GFP fusion protein were cultivated in the absence (controls) and presence of EC (200 µM) and evaluated at 9^th^ day of adulthood. DAF-16:GFP subcellular localization was classified as cytosolic, intermediate and nuclear.

**S3 Fig. Effect of EC on the expression of SOD-3 and HSP-16.2 in old worms (day 9^th^ of adult)**. Age-syncronized L1 transgenic worms of *Psod-3:gfp* and *Phsp-16.2:gfp* were cultivated in the absence (controls) and presence of EC (200 μM) in the culture media. Total GFP fluorescence of each whole worm was quantified using Image J software. Three independent experiments were performed. The results are presented as the mean values ± SEM. Approximately 35 randomly selected worms from each set of experiments were examined. Differences compared with the control (0 µM, 0.1% DMSO) were considered statistically significant at p<0.05 (*) and p<0.01 (**) and p<0.001 (***) by one-way ANOVA.

